# CRISPR metabolic screen identifies ATM and KEAP1 as targetable genetic vulnerabilities in solid tumors

**DOI:** 10.1101/2022.05.03.490553

**Authors:** Haojian Li, Yue Liu, Crystal N. Wilson, Hui Jen Bai, Maxwell Jones, Shihchun Wang, Yunjie Xiao, Jennie E. DeVore, Esther Y. Maier, Myriem Boufraqech, Urbain Weyemi

## Abstract

Cancer treatments targeting DNA repair deficiencies often encounter drug resistance, possibly due to alternative metabolic pathways that counteract the most damaging effects. To identify such alternative pathways, we screened for metabolic pathways exhibiting synthetic lethality with inhibition of the DNA damage response kinase ATM using a metabolism-centered CRISPR/Cas9 library. Our data revealed Kelch-like ECH-associated protein 1 (KEAP1) as a key factor involved in desensitizing cancer cells to ATM inhibition both *in vitro* and *in vivo*. Cells depleted of KEAP1 exhibited an aberrant overexpression of the cystine transporter SLC7A11, robustly accumulated cystine inducing disulfide stress, and became hypersensitive to ATM inhibition. These hallmarks were reversed in a reducing cellular environment indicating that disulfide stress was a crucial factor. In the TCGA pan-cancer datasets, we found that ATM levels strongly correlated with KEAP1 levels across multiple solid malignancies. Together, our results unveil ATM and KEAP1 as new targetable vulnerabilities in solid tumors.

## Introduction

Deficiencies in major DNA repair pathways are harnessed as a canonical approach to sensitize cancer cells to DNA-damaging agents. This strategy stems primarily from the notion of a synergistic relationship between two essential DNA repair pathways cooperating to mediate cancer cell survival and resistance to therapies ^1-5^. Ataxia-telangiectasia mutated (ATM) kinase controls DNA damage responses upon DNA double-strand break (DSB) formation ^6,7^, and loss of ATM kinase activity or heterozygosity for pathogenic ATM gene variants increase cancer risks. Indeed, ATM is frequently mutated across cancer types ^8^. Conversely, ATM genetic deficiency or its inhibition can be harnessed to sensitize cancer cells to radiotherapy or eliminate cancer cells with compromised DNA repair pathways. For example, ATM inhibition sensitizes lung cancer cells to Fanconi anemia (FA) pathway deficiency, a characteristic which is reflective of the shared role of ATM and FA proteins in DNA end resection during homologous recombination ^9^. A recent report demonstrated that ATM deficiency can also be utilized to sensitize metastatic breast cancer cells to inhibitors of Poly (ADP-ribose) polymerase 1 and 2 (PARP1/2), due to the synergistic roles of ATM signaling and PARP-dependent pathways in DNA repair ^10,11^. Notably, three novel ATM inhibitors (AZD0156, AZD1390, and M3541) are currently under phase I clinical trials for patients with gastric adenocarcinoma, colorectal cancer (NCT02588105), brain tumor (NCT03423628), and solid tumors (NCT03225105), potentially opening avenues for their use in combination treatments based on synthetic lethality.

Although partially effective, harnessing DNA repair deficiencies to induce an enhanced response to therapy and cell death may be hindered by the strong resistance of cancer cells to DNA-damaging agents. There is increasing evidence that ATM also controls metabolic pathways, including mitochondrial respiration, pentose phosphate pathway, and redox homeostasis, all of which enhance metastatic cancer cells survival in nutrient-poor tumor microenvironments ^12-15^. Notably, metastatic breast cancer cells rely heavily on multiple metabolic pathways, thus implying that metabolic reprogramming may be key for cancer cells to escape exposure to DNA-damaging agents ^16,17^. The pleiotropic characteristic of ATM is well-exemplified in ataxia-telangiectasia (A-T) disease, particularly with the discovery that cells lacking ATM experience oxidative stress, implying essential roles of ATM beyond its canonical DNA damage response function ^6,18^. In this regard, ATM phosphorylates heat shock protein 27 (HSP27) which in turn binds to glucose-6-phosphate dehydrogenase (G6PD), a rate-limiting enzyme of the pentose phosphate pathway (PPP), to promote the synthesis of NADPH and ribose-5-phosphate^12^. Elevated NADPH increases antioxidant capacity while ribose-5-phosphate functions as a precursor for *de novo* nucleotide synthesis. ATM also promotes PPP by up-regulating transcription of G6PD gene in response to increased mitochondrial ROS ^19-21^. Furthermore, ROS are also produced from peroxisomes and NADPH oxidases. ATM regulates the turnover of peroxisomes by inducing pexophagy in response to ROS ^22^. We also demonstrated that the NADPH oxidase 4 (NOX4) is up-regulated in A-T cells and in normal cells treated with ATM inhibitor ^13^. These findings suggest roles for ATM in reducing ROS via pexophagy or down-regulation of NADPH oxidases. A recent report from Herrup’s group demonstrated that ATM controls mitochondrial biogenesis by phosphorylating the Nuclear Respiratory Factor 1 (NRF1) in the brain ^23^. In line with these evident links between ATM and metabolism, we demonstrated that deletion of histone H2AX, a major substrate of ATM kinase, impairs mitochondrial function and promotes repression of the mitochondrial biogenesis activator, PGC1-α ^24^. However, the interplay between ATM and metabolism, and the clinical relevance of this link in cancer is yet to be investigated, highlighting major gaps in our understanding of how ATM and other DNA repair genes control various metabolic pathways in cancer cells.

In this study, we utilized a metabolism-centered CRISPR screening to evaluate the metabolic reprogramming ability of cancer cells upon resistance to ATM inhibition. We identified Kelch-like ECH-associated protein 1 (KEAP1), an oxidative stress sensor that controls the Nrf2-regulated antioxidant pathway, as a critical driver for resistance to ATM kinase inhibition. KEAP1 depletion endows cancer cells with a strong ability to heavily rely on cystine metabolism. This effect is mediated by an aberrant expression of NRF2-regulated cystine transporter SLC7A11. Inhibition of disulfide stress rescues cell death promoted by a concomitant elevation in cystine uptake and NADPH depletion resulting from KEAP1 loss and ATM inhibition, respectively. Remarkably, pharmacological inhibition of ATM abrogates the growth of KEAP1-deficient breast and lung tumors, which is commonly observed in human cancers. Our data reveal that ATM levels strongly correlate with KEAP1 levels across multiple solid malignancies, and establish ATM and KEAP1 as vulnerabilities in solid tumors that can be targeted therapeutically.

## Results and discussion

### Identification of KEAP1 as a critical gene for resistance to ATM inhibition

To identify metabolic genes whose loss enhances cell killing by ATM inhibitors, we first evaluated the efficacy of the ATM inhibitor AZD1390 using triple-negative breast cancer cells MDA-MB-231 and hydrogen peroxide as an ATM activator. Low doses of AZD1390 in the nanomolar range, abolished the activated form of ATM over 1 h post-drug treatment (Figure 1a). Using proliferation assays, we evaluated the degree to which AZD1390 promotes cell death. As expected, only up to 30% of cell death was observed 8 days following exposure to doses equivalent to or higher than 30 nM of AZD1390 (Figure 1b). These data imply a considerable resistance to ATM inhibitor when used as a single agent. We performed a CRISPR-based negative selection screen for metabolic genes whose loss enhances cell killing by ATM inhibitors (Figure 1c). We surmise that such genes would endow cancer cells with an elevated ability to resist and survive upon ATM inhibition. To selectively evaluate the contribution of metabolism to such resistance, we utilized a previously established metabolism-centered CRISPR/Cas9 library^25^, which consists of ∼30,000 sgRNA targeting ∼3,000 metabolic enzymes, small molecules transporters, and metabolism-related transcription factors (∼10 sgRNA/gene), and 500 control sgRNAs in a Cas9-expressing lentiviral vector. MDA-MB-231 cells were infected with a low multiplicity of infection (MOI = 0.3) to deliver less than one virus particle per cell to ensure effective barcoding of individual cells. Following a week of selection, cells were either treated with a moderate dose of AZD1390 (50 nM) or with DMSO for 14 days. Cells were collected pre- and post-drug selection, genomic DNA was extracted, and sgRNA sequences were PCR amplified. Using massive parallel sequencing, we measured the abundance of all the sgRNAs in the vehicle and ADZ1390-treated cells at the beginning and the end of the culture period. To assess the degree to which genes are required for survival upon ATM inhibition, we calculated for each gene the beta-score using the *MAGeCK MLE* pipeline. The beta-score for a specific gene is the change in the abundance of the 10 sgRNAs targeting the gene when comparing the drug-treated group to the Day 0 untreated group. Our data revealed Aryl Hydrocarbon Receptor Nuclear Translocator (ARNT) as potentially the most required gene for survival upon ATM inhibition (Figure 1d, e). However, the beta scores for each of the 10 sgRNA guides for ARNT indicate that most guides score positively in both DMSO and AZD1390-treated specimens (Supplementary Figure 1a), implying that ARNT deletion may only alter cell proliferation, rather than promote sensitivity to ATM inhibition. Consistent with these observations, 9 of the 10 sgRNAs for Kelch-like ECH-associated protein 1 (KEAP1) gene, the second most required hit, show negative beta-scores in cells treated with AZD1390, among them 5 sgRNAs show positive beta-scores in cells treated with DMSO (Supplementary Figure 1b). A similar pattern of distribution was observed for Phosphate And Tensin Homolog (PTEN) gene (Supplementary Figure 1c). Indeed, PTEN loss sensitizes colorectal cancer cells to ATM inhibitors ^26^. Therefore, we surmise that KEAP1 is an important gene required for survival upon ATM inhibition.

**Figure 1.**
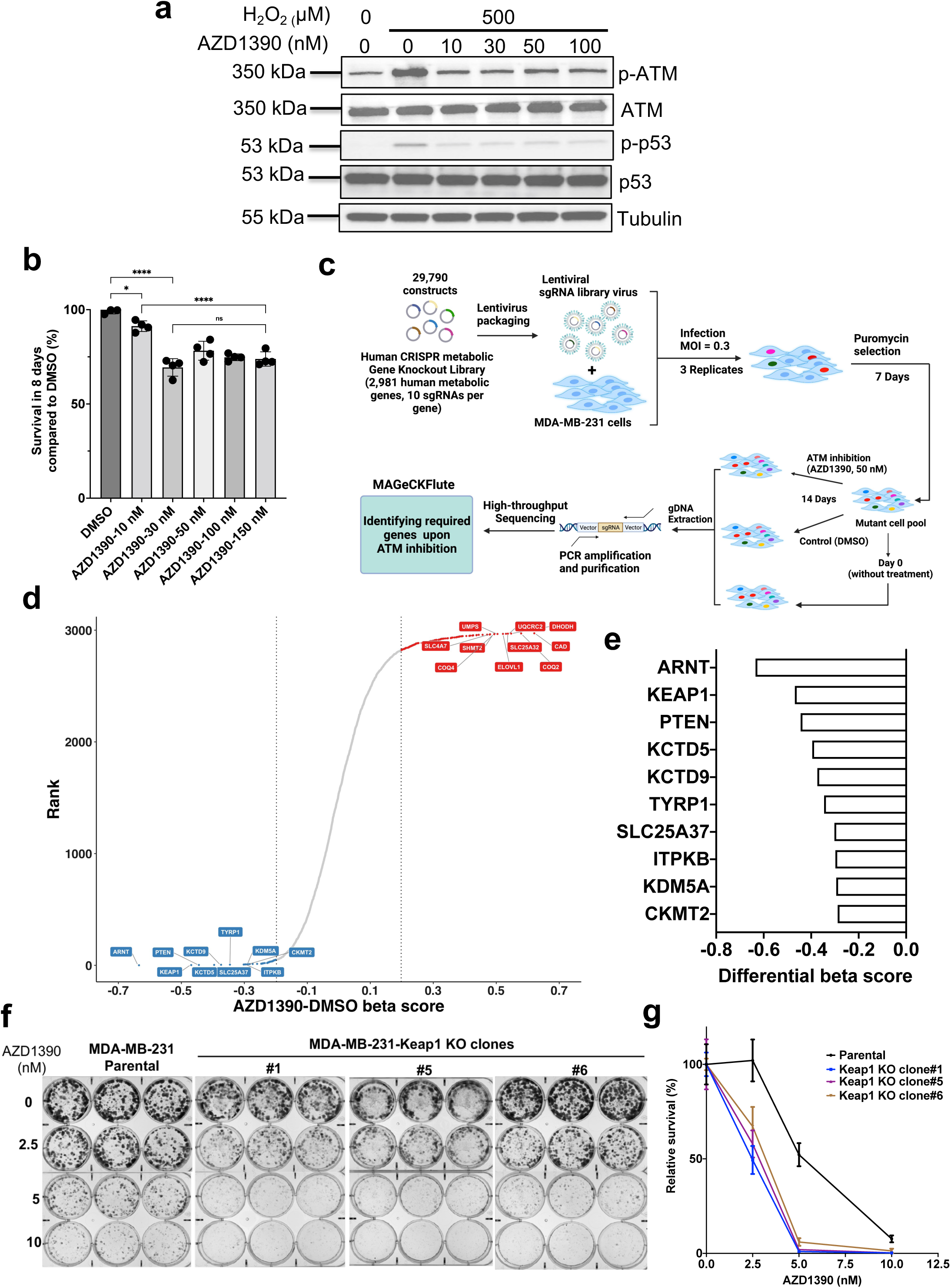
Metabolism-centered CRISPR/Cas9 knockout library screen identified KEAP1 as a driver for resistance to ATM inhibition. (a) ATM kinase activity inhibition by AZD1390. MDA-MB-231 cells were pre-treated with the ATM inhibitor AZD1390 as indicated (10, 30, 50, 100 nM) for 1 h followed by the addition of H_2_O_2_ (500 μM) for 30 min. Phospho-p53-Ser^15^ (p-p53) and phospho-ATM-Ser^1981^ (p-ATM) induced by H_2_O_2_ were probed by western blot and analyzed for ATM kinase activity. Levels of p53, ATM and tubulin were used as a control. (b) Survival of MDA-MB-231 cells after treatment with indicated AZD1390 concentrations for 8 days. Statistical significance was determined by one-way ANOVA with Tukey’s multiple comparisons test. Data are represented as mean ± SD; n = 4. *, p < 0.05; ****, p < 0.0001. ns, not significant (c) Schematic diagram illustrating the workflow of metabolism-centered CRISPR/Cas9 knockout library screen. (d) The Rank plot of genes generated by MAGeCKFlute, which is sorted based on differential beta score by subtracting the DMSO beta score from the AZD1390 beta score. (e) The top ten genes with the lowest differential beta score. (f) Clonogenic assays of parental and *KEAP1* knockout cells (clones #1, #5, #6) after treatment with ATM inhibitor (AZD1390) as indicated (2.5, 5, 10 nM). (g) Quantification of panel f.

To validate the results from our CRISPR/Cas9 knockout library screen, we generated both ARNT and KEAP1 knockout clones by infecting MDA-MB-231 cells with sgRNAs included in the library. We measured cell survival upon ATM inhibition using clonogenic assays. Our data revealed that KEAP1 loss robustly sensitizes cancer cells to both AZD1390 (Figure 1f, g) and the canonical ATM inhibitor KU60019 (Supplementary Figure 2 a-c). Specifically, while 50% of MDA-MB-231 parental cells remain viable upon treatment with 5 nM of AZD1390, only ∼ 1% of KEAP1-knockout cells survived 12 days post drug treatment. Inversely, when exposed to ATM inhibitors AZD0156 and AZD1390, ARNT knockout (ARNT KO) cells exhibit a similar degree of sensitivity as compared to parental cells (Supplementary Figure 3 a-e), confirming that ARNT loss is not synthetically lethal with ATM inhibition. To ascertain a requirement for both ATM and KEAP1 in cell survival in another model of breast cancer, we silenced KEAP1 expression in luminal breast cancer cell line MCF7 using small interference RNAs (siRNAs). Following exposure to ATM inhibitor AZD1390, KEAP1-deficient MCF7 cells were more sensitive to ATM inhibition than control cells, observations which are consistent with results obtained in KEAP1-KO MDA-MB-231 cells (Supplementary Figure 4a, b). These findings confirm that KEAP1 loss is synthetically lethal with ATM inhibition. To further determine whether cooperation between KEAP1 and ATM is essential for resistance, MDA-MB-231 cells were treated with the combination of AZD1390 and dimethyl fumarate (DMF), an FDA-approved KEAP1-NRF2 interaction inhibitor ^27,28^. Measurement of cell death in clonogenic assays revealed over 85% of cell death following a concomitant inhibition of KEAP1 and ATM, while only ∼30% of cell death was detected with single agents treatment (Supplementary Figure 4 c, d). Collectively, these results reveal a previously unknown synergistic relationship between ATM and KEAP1 in breast cancer cells. While most available KEAP1 inhibitors or molecules targeting KEAP1-NRF2 interaction have yet to advance beyond preclinical cancer models presumably due to their lower selectivity, our findings provide unique evidence of concomitantly targeting a metabolic pathway and DNA repair genes to promote synthetic lethality in cancer cells.

### ATM correlates with KEAP1 across human cancers

KEAP1 and ATM are frequently mutated across cancer types ^8,29^. To investigate whether the synergistic relationship between KEAP1 and ATM may provide clues for any clinical correlation, we conducted an expression level comparison of KEAP1 and ATM in a large panel of human malignancies including a cohort of solid tumors of multiple origins (pan-cancers cohort), a cohort of invasive breast cancer patients, and a cohort of patients with lung adenocarcinoma from The Cancer Genome Atlas (*TCGA*) datasets. The data indicate that the transcript levels of ATM and KEAP1 inversely correlated across a large panel of over 10, 071 specimens from multiple origins (Figure 2a), in specimens from breast invasive carcinoma (Figure 2b), as well as in lung adenocarcinoma (Figure 2c), as revealed by the Spearman correlation coefficients (pan-cancers panel, n = 10, 071, r = -0.35, p < 2.2e^-16^; breast invasive carcinoma panel, n= 994, r = -0.4, p < 2.2e^-16^, lung adenocarcinoma, n= 503, r = -0.24, p = 3.9e^-08^). Collectively, these observations suggest that at least in a subset of patients, KEAP1 and ATM may synergistically cooperate to inhibit cancer cell growth, providing unique clues that vulnerability in both KEAP1 and ATM can be harnessed to increase cancer cell response to therapy.

**Figure 2.**
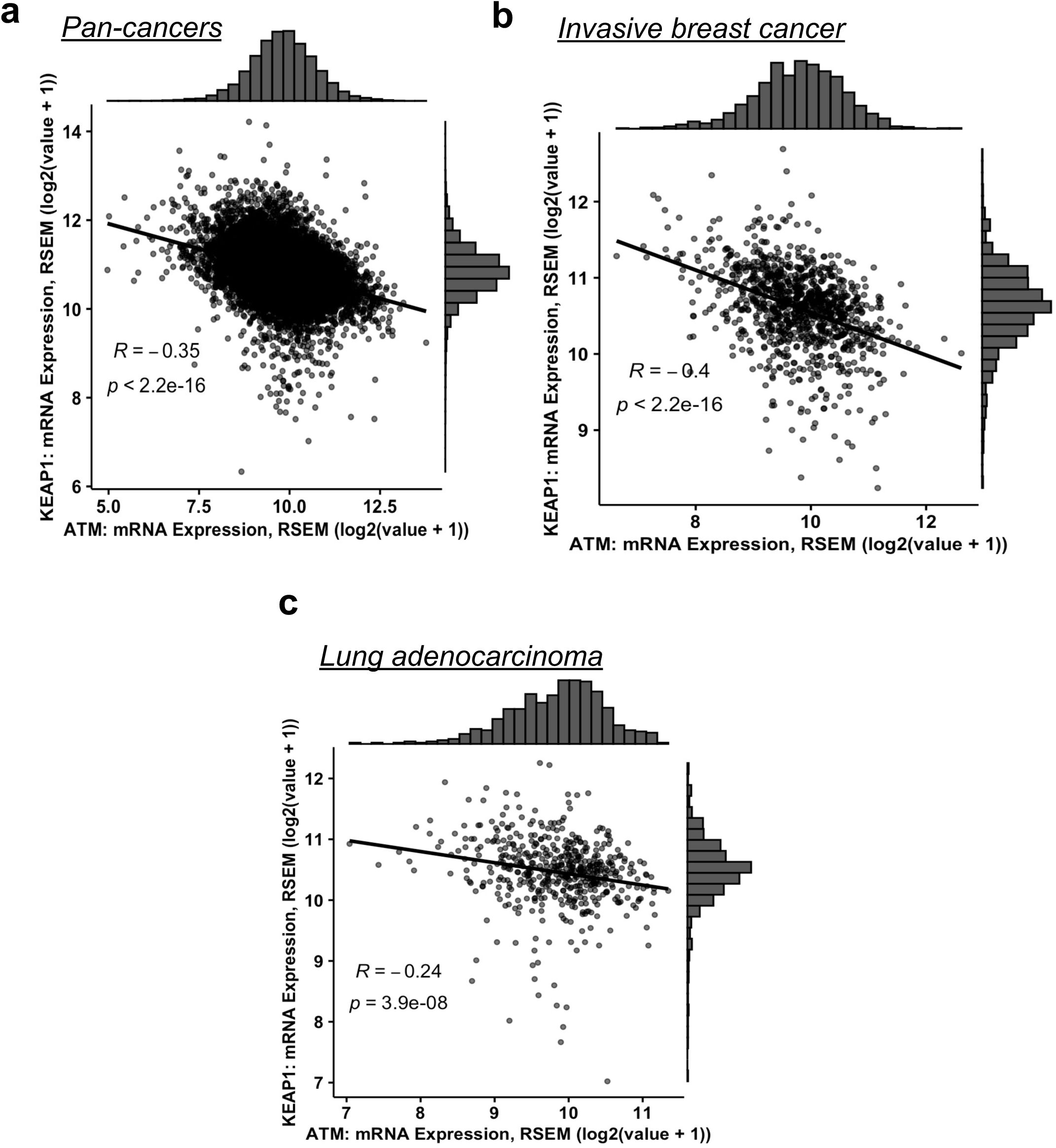
Statistically significant correlation between transcript levels for ATM and KEAP1 across cancers. (a) 10,071 pan-cancer samples from multiple malignancies in the TCGA datasets. (b) 994 invasive breast cancer samples in the TCGA dataset. (c) 503 lung adenocarcinoma samples in the TCGA dataset. Spearman r and p value is shown in the panel. The Log2 expression level is shown for the indicated genes.

### KEAP1 deficiency sensitizes breast and lung tumors to ATM inhibition

To determine whether KEAP1 is essential for resistance to the pharmacological inhibition of ATM *in vivo*, we generated cancer cells xenografts in immunocompromised *NOD SCID gamma* (NGS) mice utilizing both parental and KEAP1 knockout MDA-MB-231 metastatic breast cancer cells. When the tumors became visible in bioluminescence-based imaging post inoculation, mice were randomized and treated daily with either an ATM inhibitor (AZD0156 or AZD1390) or with a vehicle (Figure 3a). Treatment with AZD0156 led to an ∼85% attenuation of tumor growth when KEAP1 was deleted (Figure 3b, c), compared to parental tumor cells which grew steadily. Parental tumor cell growth was partly impaired when mice were treated with AZD0156 alone, and a similar pattern of tumor growth was observed with KEAP1-deleted cells treated with vehicle only. Although very moderate and non significant, the effect of ATM inhibitor when used as a single agent, partially recapitulates previous findings that ATM silencing is sufficient to reduce triple-negative breast cancer tumors progression in the lung ^30^. Remarkably, the elevated sensitivity of KEAP1-deficient cells to ATM inhibitor was consistent with a 2-fold increase in median survival post AZD0156 treatment (18 days versus 37.5 days) (Figure 3d). More noticeably, we showed that AZD0156 treatment did not yield adverse effects, nor changes in body weight throughout the treatment (Supplementary Figure 5), implying that the drug is relatively well tolerated in animals. To confirm that our study was not limited to the use of one single ATM inhibitor, we treated mice with AZD1390, the ATM inhibitor used in our CRISPR/Cas9 knockout library screening. While the treatment with AZD1390 alone marginally reduced MDA-MB-231 tumor growth in the lung, KEAP1 loss remarkably enhanced tumor sensitivity to ATM inhibition (Figure 3e, f). The number and the size of lung nodules remain noticeably high when mice were treated with AZD1390 alone, while KEAP1 loss led to a drastic reduction in lung nodules in mice treated with AZD1390, as revealed by the relatively healthy appearance of the lung (Figure 3g, h). These findings suggest that inactivation of KEAP1 is sufficient to sensitize breast tumors to the pharmacological inhibition of ATM *in vivo*, strongly raising the possibility of combining inhibition of DNA repair genes with selective repression of essential metabolic genes to enhance response to therapy.

**Figure 3.**
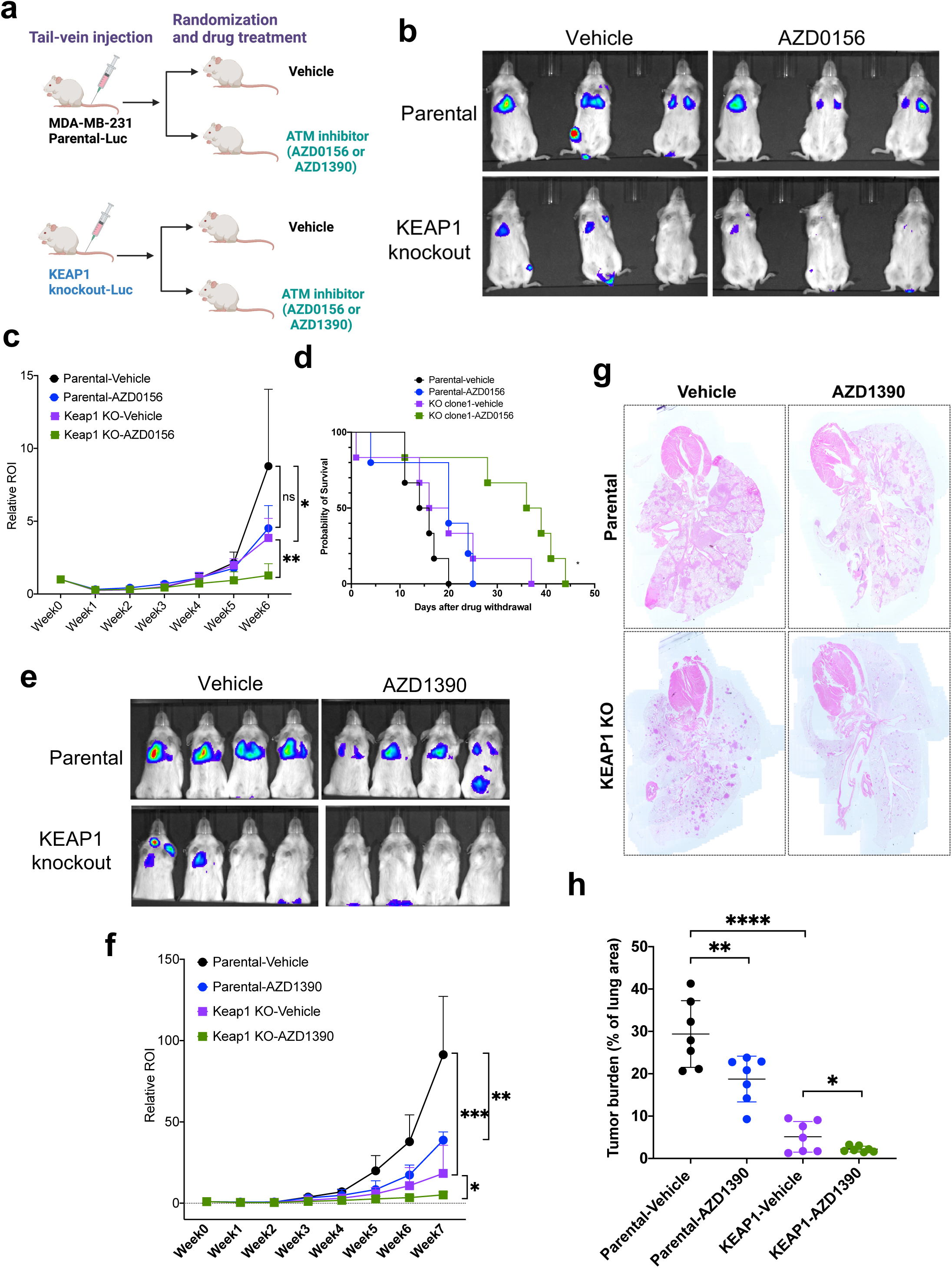
Depletion of KEAP1 sensitized triple-negative breast cancer cells MDA-MB-231 to ATM inhibition in immunocompromised NOD SCID gamma mice. (a) Schematics of the experimental metastasis model using MDA-MB-231-luciferase cells. (b) Representative bioluminescence imaging of mice injected with MAD-MB-231-Luciferase parental or KEAP1 KO clone #1 and treated with vehicle or ATM inhibitor (AZD0156, 20 mg/kg) at week 6. (c) Weekly tumor progression in mice with indicated treatment based on the bioluminescence imaging by IVIS system. Statistical significance was determined by one tail unpaired welch’s t test. Data are represented as mean ± SD; n = 5 or 6. *, p < 0.05; **, p < 0.01. (d) Survival of mice following the end of treatment with AZD0156 Statistical significance was determined by log-rank test. * indicates p<0.05 comparing KO clone1-vehicle (purple curve) to KO clone1-AZD0156 (green curve). (e) Representative bioluminescence imaging of mice injected with MDA-MB-231-Luciferase parental or KEAP1 KO clone #1 treated with vehicle or ATM inhibitor (AZD1390, 20 mg/kg) at week 7. (f) Weekly tumor progression in mice with indicated treatment based on the bioluminescence imaging by IVIS system. (g) Representative hematoxylin and eosin (H&E) staining using heart and lung tissue from mice in (f) harvested at week 7. Note that cells with depleted KEAP1 exhibited strikingly elevated numbers of nodules, most of which are particularly smaller compared to parental tumors (black arrows). (h) Quantification of area occupied by tumor cells using hematoxylin and eosin (H&E) staining of lung from mice in (e) harvested at week 7 (mean ± SD, n = 7). Statistical significance was determined by with one tail unpaired welch’s t test. Data are represented as mean ± SD; n = 7. *, p < 0.05; **, p < 0.01; ***, p < 0.001; ****, p < 0.0001.

KEAP1 is mutated in about 3.93% of all cancers, with the highest frequency of mutation in non-small cell lung cancer (NSCLC) at 15.8% ^31-33^. We surmise that KEAP1 mutation, its inactivation or repression can therefore be harnessed to sensitize lung cancer cells to synergistic players, with the ATM kinase identified in this study as a strong candidate. To evaluate the relevance of the synergistic relationship between ATM and KEAP1 in lung tumors, we analyzed the effect of the pharmacological inhibition of ATM in KEAP1-deficient lung cancer cells. Specifically, we tested AZD1390 in xenograft models established from control and KEAP1-depleted cells generated using short hairpin RNAs (shRNAs) interference approach (Figure 4a). Both control and KEAP1-depleted cells were transfected with a luciferase-expressing construct to assess tumor growth *in vivo* using bioluminescence-based imaging (Figure 4b). Our data revealed that repression of KEAP1 alone in NCI-H1299 significantly reduces tumor growth, findings which may not reflect the tumor-promoting function of KEAP1 deficiency in the lung (Figure 4c-e). Pharmacological inhibition of ATM with AZD1390 did not yield a noticeable effect on tumor growth in mice injected with NCI-H1299 sh-control cells. However most strikingly, KEAP1 depletion drastically sensitizes lung cancer cells to ATM inhibition (Figure 4c, e). This effect was substantiated by the dramatic reduction in tumor volume and tumor weight at the endpoint (Figure 4f, g). Collectively, these data validate the synthetic lethality between KEAP1 loss and ATM inhibition in lung cancer cells, findings that strongly recapitulate observations with metastatic breast cancer cells. While the reduction in tumor growth observed in KEAP1-deficient cells may not substantiate the tumor-promoting role of KEAP1 mutations, particularly in lung cancers, KEAP1-deficient cells may still retain their highly invasive phenotype. Nonetheless, our study uniquely unveils the extent to which KEAP1 deficiency can be harnessed to sensitize tumor cells to ATM inhibition, opening avenues of research involving ATM and KEAP1 as novel therapeutic targets.

**Figure 4.**
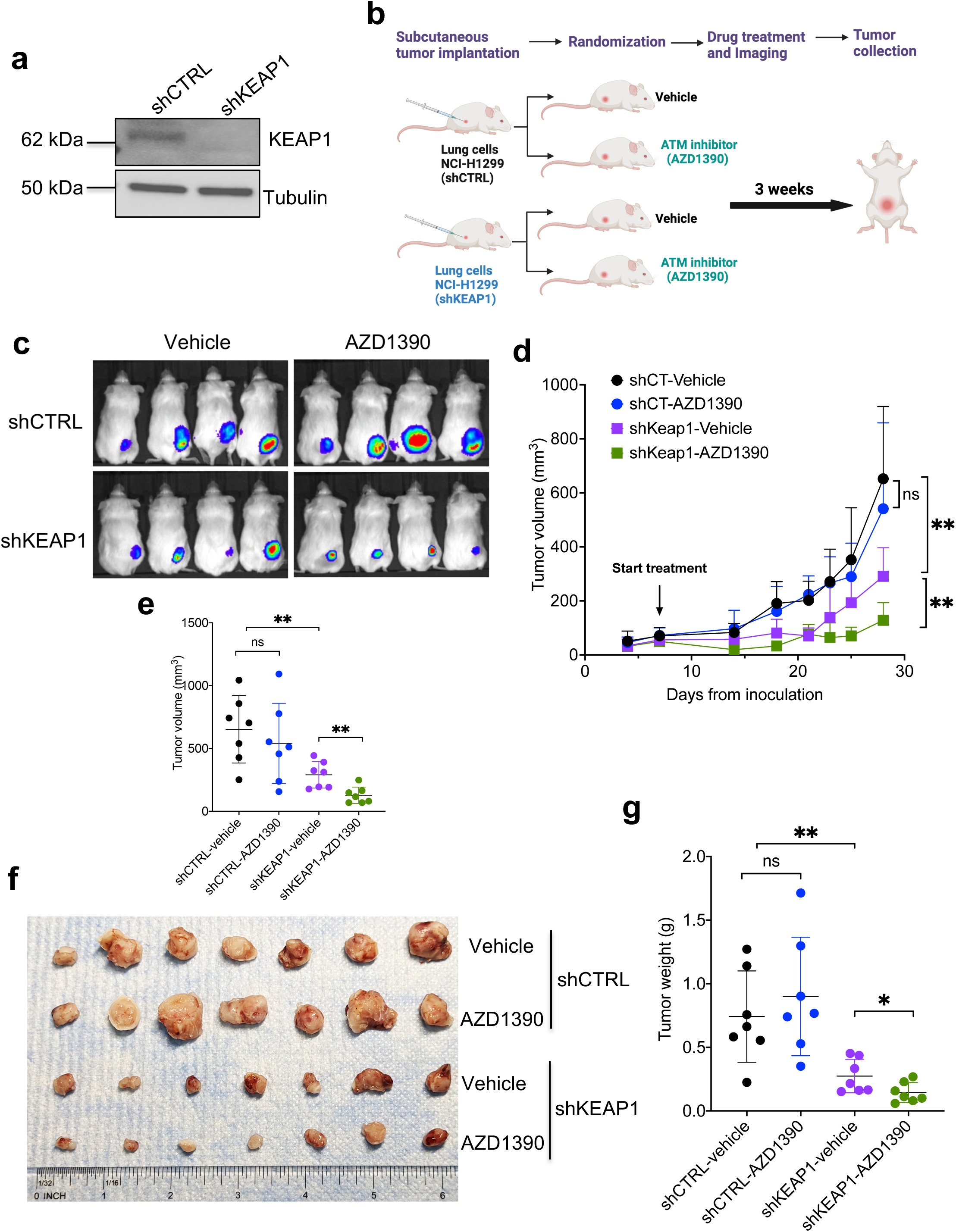
Knockdown of KEAP1 sensitized non-small lung cancer cells NCI-H1299 cells to ATM inhibition in immunocompromised NOD SCID gamma mice. (a) Western blot showing *KEAP1* knockdown level in NCI-H1299-luciferase cells. (b) Schematics of lung cancer xenograft model using NCI-H1299-luciferase cells. (c) Representative bioluminescence imaging of mice inoculated with NCI-H1299-Luc shcontrol (shCTRL) or shKEAP1 cells treated with vehicle or ATM inhibitor (AZD1390, 20 mg/kg) at week 4. (d) Measurement of tumor volume throughout the experiment. (e) Tumor volume at week 4. (f) Image of tumors taken *ex vivo* at week 4. (g) *Ex vivo* tumor weight at week 4. Statistical significance was determined by two-tail unpaired student t test. Data are represented as mean ± SD; n = 7. *, p < 0.05; **, p < 0.01.

### Cell death by ATM inhibition in KEAP1-depleted cells is partly caused by aberrant disulfide bonds buildup and stress upon NADPH depletion

KEAP1 is a substrate adaptor for an E3-ubiquitin ligase complex with a high binding affinity for nuclear factor erythroid 2-related factor 2 (NRF2). KEAP1 controls redox homeostasis via its dissociation from NRF2, followed by NRF2 nuclear translocation, and subsequent activation of NRF2-regulated antioxidant response genes ^34,35^. KEAP1 loss leads to high dependency on glucose metabolism ^33^. A striking recent report by Gan’s group established that, under glucose starvation, KEAP1 deficiency causes an elevated cystine uptake via aberrant overexpression of the NRF2-regulated cystine transporter SLC7A11, followed by accumulation of disulfide molecules, with the oxidized glutathione (GSSG) being one of the most abundant forms ^33^. The findings provided a piece of strong evidence that glucose deprivation coupled with KEAP1 loss may be responsible for the toxic accumulation of disulfide molecules upon NADPH pool depletion, a phenomenon described as disulfide stress. The buildup of disulfide molecules is characteristic of a profound depletion in reduced glutathione (GSH) pool, subsequently leading to redox system collapse and cell death. Indeed, cancer cells depend heavily on GSH synthesis primarily from the cysteine pool to overcome the high demand for antioxidant defense required for survival and resistance to therapy ^36-38^. Consistent with the model of glucose starvation in KEAP1-deficient cells ^33^, we surmise that NADPH depletion by ATM inhibition may synergize with KEAP1 loss to promote the accumulation of disulfide molecules and cytotoxicity due to depletion in the glutathione pool. As expected, KEAP1 loss led to a robust accumulation of NRF2 and overexpression of its target SLC7A11 (Figure 5a). These observations raise the possibility that KEAP1-deficient cells may heavily rely on cystine uptake for GSH synthesis and survival upon stress. As such, NADPH depletion by ATM inhibition may drastically impair the survival of KEAP1-deficient cells, presumably via disulfide molecules buildup, and an imbalance in GSH/GSSG ratio. More specifically, depletion in the pool of GSH may favor an environment prone to accumulation of disulfide molecules such as GSSG, and disulfide stress. Indeed, previous studies demonstrated that ATM increases the pool of NADPH by promoting the activity of glucose-6-phosphate dehydrogenase (G6PD), a rate-limiting enzyme of the pentose phosphate pathway (PPP) ^12^. To determine the degree to which ATM may impact NADPH production, we measured the NAPDH pool in parental and KEAP1-deficient cells upon ATM inhibition. As shown in Figure 5b, ATM inhibition is accompanied by a significant depletion in the NADPH pool 14 days post drug treatment. Most interestingly, these observations are paralleled by a drastic reduction in the GSH/GSSG ratio in KEAP1 knockout cells from 6 days up to 14 days post drug treatment (Figure 5c, d). Strikingly, ATM inhibition strongly potentiates these effects.

**Figure 5.**
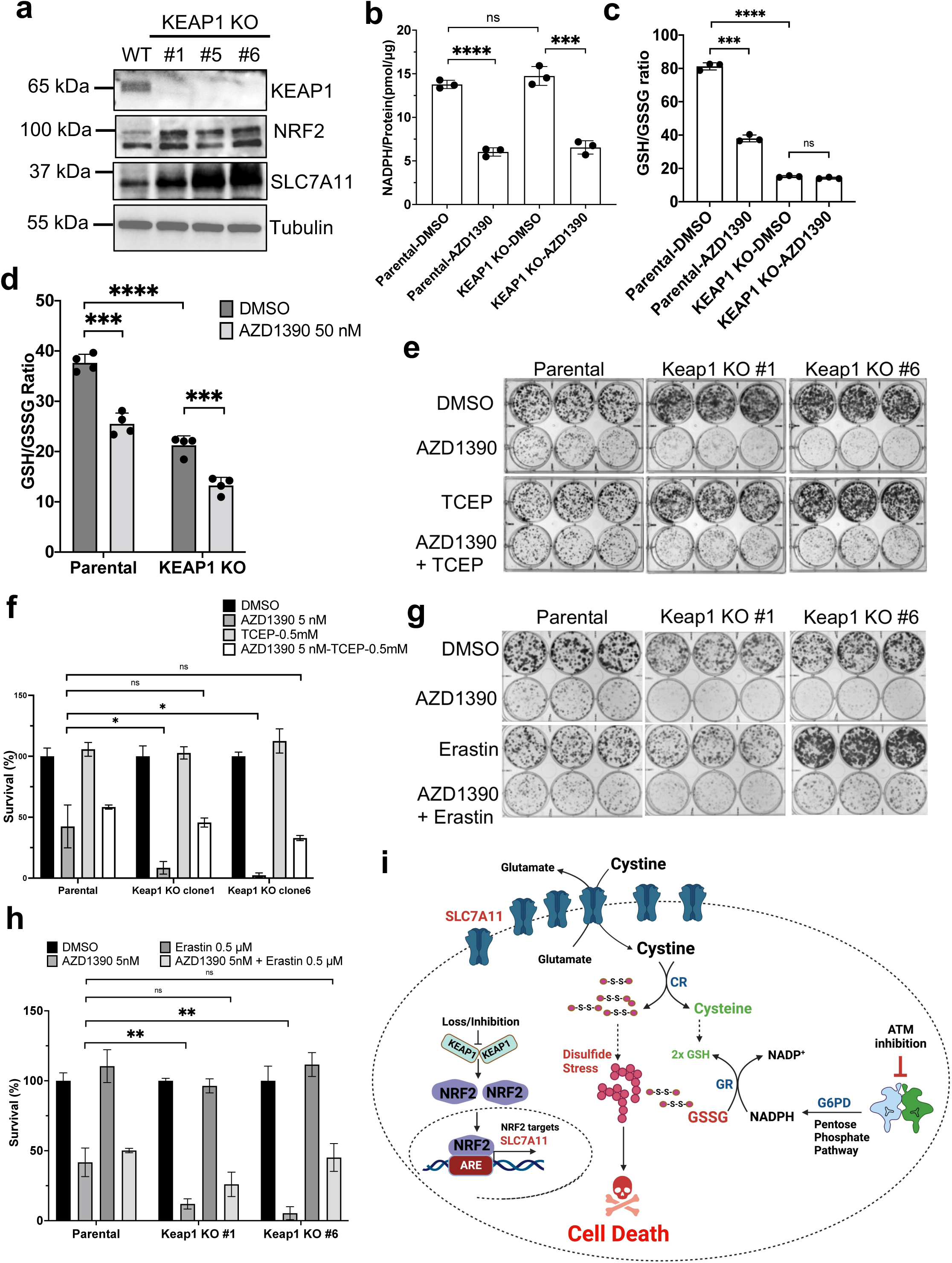
Cell death by ATM inhibition upon KEAP1 loss is partly caused by aberrant disulfide bonds buildup and stress upon NADPH depletion. (a) Activation of *SLC7A11* in KEAP1-deficient cells. NRF2 and SLC7A11 levels were probed by western blot using MDA-MB-231-Luc parental, KEAP1 KO clone#1, 5 and 6. Tubulin was used as a loading control. (b) Concentrations of NADPH in MDA-MBA-23-luc cells upon ATM inhibition for 14 days. (c-d) Ratio of GSH/GSSG in MDA-MBA-23-luc cells upon ATM inhibition for 6 days (c) or for 14 days (d). (e) Clonogenic assays of parental and *KEAP1* knockout cells (clones #1, #6) after treatment with ATM inhibitor (AZD1390) supplemented with TCEP at 0.5 mM. (f) Quantification of (e). (g) Clonogenic assays of parental and *KEAP1* knockout cells (clones #1, #6) after treatment with ATM inhibitor (AZD1390) supplemented with Erastin at 0.5 μM. (h) Quantification of (g). (i) Model for cell death induced by aberrant disulfide bonds buildup in KEAP1-deficient cells upon ATM inhibition. Statistical significance was determined by two-tail unpaired student t test. Data are represented as mean ± SD; n = 3 or 4. *, p < 0.05; **, p < 0.01; ***, p < 0.001; ****, p < 0.0001; ns, not significant.

To determine whether reduction of disulfide molecules or inhibition of cystine transporter (SLC7A11) is sufficient to rescue KEAP1-deficient cells from death upon ATM inhibition, we treated parental and KEAP1 knockout cells with the disulfide molecules reducing agent tris-(2-carboxyethyl)-phosphine (TCEP), or with Erastin respectively. Remarkably, the TCEP treatment rescued cell death induced by ATM inhibition in KEAP1-deficient cells to the level of AZD1390-treated parental cells (Figure 5e, f). Likewise, treatment with Erastin led to a similar pattern of mitigation (Figure 5g, h), further supporting that disulfide molecules buildup upon ATM inhibition strongly sensitizes KEAP1-deficient cells to death. Collectively, these findings suggest that KEAP1-deficient cells heavily rely on cystine metabolism to produce sufficient amounts of reduced glutathione (GSH) required for their survival in a highly antioxidant system-demanding environment, and that depletion of NADPH upon ATM inhibition overloads cells with disulfide molecules, thus leading to cell death (Figure 5i).

The aberrant overexpression of SLC7A11 in KEAP1-depleted cells implies an elevated potential for cystine accumulation. While our current study does not provide evidence of cystine accumulation in KEAP1-deficient cells, it should be noted that the cystine molecules directly uptaken by SLC7A11 at the cell membrane are likely rapidly converted into cysteine due to the existence of cystine reductase (CR). Inversely, a lack of sufficient amount of other reductases responsible for converting disulfide molecules into their reduced forms may underlie the lower GSH/GSSG ratio observed in KEAP1-depleted cells, particularly in the absence of a functional ATM (Figure 5i).

We have identified KEAP1 and ATM as novel targetable vulnerabilities in breast and lung cancers utilizing metabolism-centered CRISPR/Cas9 screening. Our study provides strong evidence supporting the stratification of patients with KEAP1-mutant or NRF2-hyperactivated tumors as likely responders to ATM inhibitors. We showed that KEAP1 is particularly required for invasive breast and lung cancer cells to survive upon ATM inhibition. Beyond targeting KEAP1-mutant lung cancers, our study opens up a unique line of investigation on using the combination of ATM and KEAP1 inhibitors in multiple human malignancies to yield an enhanced response to therapy.

The significance of the findings in this study lies in two major areas. One is the identification of an unprecedented synthetic lethality between inhibition of the DNA repair response kinase ATM and deficiency in KEAP1, a master regulator of redox homeostasis in mammalian cells. KEAP1 is mutated in about 3.93% of all cancers, with the highest frequency of mutations in non-small cell lung cancer (NSCLC) at 15.8%. Patients with NSCLC exhibit a significantly lower median survival, owing to the lack of effective therapeutic options and the immune-evasive phenotype of KEAP1-mutant cancer cells. The second major significance of our study is that we provide unique clues for stratifying patients with KEAP1-mutant or NRF2-hyperactivated tumors as likely responders to ATM inhibitors.

## Materials and Methods

### Compounds

ATM inhibitors AZD1390 and AZD0156 were generously provided by *AstraZeneca via* a Materials Transfer Agreement (MTA). Both drugs were pharmaceutical grade, and were used for the *in vivo* studies. All *in vitro* experiments were conducted using AZD1390 (Cat. S8680), AZD0156 (Cat. S8375), Ku-60019 (Cat. S1570), Erastin (Cat. S7242) from *Sellechchem*. Dimethyl fumarate (DMF) (Cat. 242926-100G) and Tris(2-carboxyethyl)phosphine hydrochloride (TCEP) (Cat. C4706-10G) were purchased from Sigma-Aldrich.

### Metabolism-centered CRISPR/Cas9 knockout library screen

In this study, the human CRISPR metabolic gene library was used to identify metabolic genes responsible for ATM inhibition resistance in the breast cancer cell line, MDA-MB-231. The library was a gift from David Sabatini’s laboratory to Addgene (Addgene #110066). The CRISPR knockout library screen was performed as described previously ^39^. Briefly, we transduced the library which contains 29,790 gRNAs targeting 2981 human metabolic genes (∼10 gRNAs per gene and 500 control gRNAs targeting intergenic region) at a low MOI (∼0.3) to ensure effective barcoding of individual cells. Then, the transduced cells were selected with 1 μg ml^-1^ of puromycin for 7 days to generate a mutant cell pool, which were then split into 3 groups, one group was then frozen and designated as Day 0 sample, the other two groups were treated with vehicle (DMSO) and AZD1390 (50 nM) for 14 days, respectively. After treatment, at least 16 million cells were collected for genomic DNA extraction to ensure over 500X coverage of the human CRISPR metabolic gene library. The sgRNA sequences were amplified using NEBNext^®^ High-Fidelity 2X PCR Master Mix and subjected to Next Generation Sequencing by the Genomic Sequencing and Analysis Facility of the University of Texas at Austin. The sgRNA read count and hits calling were analyzed by MAGeCK^40^.

### Cell culture and plasmid transient transfection

The human breast cancer cell lines MDA-MB-231 (from ATCC, Manassas, VA USA) and MCF7 (from ATCC, Manassas, VA USA), were grown at 37°C with 5% CO_2_ in DMEM (Fisher Scientific, Waltham, USA), supplemented with 10% Fetal Bovine Serum (Thermo Scientific, Waltham, USA). Cells were authenticated using *a colorimetric signal amplification system*, and tested for mycoplasma contamination (R&D systems, Minneapolis, USA). All media were supplemented with penicillin and streptomycin (Gibco, Amarillo, USA).

### Generation of stable luciferase expression MDA-MB-231 (231-Luc) cells

For lentiviral infection, HEK-293 FT cells were used for virus package according to the manufacturer’s instructions. To obtain luciferase expression cells (231-Luc), MDA-MB-231 parental cells infected with pF-CAG luc hygro (Addgene #67502) were selected with hygromycin (400 μg ml^-1^) for 1 week and the expression of luciferase was examined by IVIS lumina instrument. Briefly, the lentivirus packaging system was used as follows: lentiviral construct that expressed luciferase was co-transfected with lentiviral packaging plasmids psPAX2 and pCMV-VSVG into HEK-293 FT cells using Lipofectamine® 3000 (Thermo Scientific, Waltham, USA) according to the manufacturer’s instructions. At 48 hours post-transfection, the culture medium was collected to be incubated with MDA-MB-231 cells in the presence of polybrene (8 μg. ml^-1^). At 72 hours post-infection, infected cells were selected with hygromycin B (400 μg ml^-1^) to establish a stable cell line.

### Establishment of knockout and knockdown cell lines

To generate MDA-MB-231-Luc cells expressing the indicated sgRNAs, the oligonucleotides described below were cloned into the pLentiCRISPR V2 viral vector (Addgene, # 52961). sgKEAP1: 5’-GGGCCGCCTGATCTACACCG - 3’. sgARNT: 5’-GCAATGGCTGGACCCAGTGA - 3’. To generate KEAP1 knockdown stable model, NCI-H1299 cells were infected with KEAP1 targeting shRNA expressing plasmids (pLKO.1) lentiviral particles. The shRNA sequence used is listed as follows: shKEAP1: 5’-GTGGCGAATGATCACAGCAAT - 3’.

### siRNA transfection

For siRNA transfection, 0.3 million cells were harvested and reverse transfected with a specific siRNA duplex at 25 nM using Lipofectamine RNAiMAX Reagent (Invitrogen, Cat. No.13778075) according to the manufacturer’s instructions for 48 hr. The siKEAP1 (4392420; Assay ID: s18981) and negative control siRNA (4390843) were obtained from Ambion™.

### Western blots

Cells were washed twice with PBS (Thermo Scientific, Waltham, USA), directly solubilized in RIPA buffer, and then subjected to SDS-PAGE. Proteins were electro-transferred to 0.2 µm nitrocellulose sheets (Bio-Rad, Hercules, USA) for immunodetection with the following primary antibodies: ATM (1:1000; 2873S, Cell Signaling Technology Inc, MA, USA); Phospho-ATM (1:1000; 13050S, Cell Signaling Technology Inc); p53 (1:1000; GTX70214, GeneTex, CA, USA); Phospho-p53 (1:1000; 9286S, Cell Signaling Technology Inc); Tubulin (1:2500; T9026-.2ML, Sigma-Aldrich, MO, USA); NRF2 (1:1000; 12721S, Cell Signaling Technology Inc); GAPDH (1:2500; 3683S; Cell Signaling Technology Inc, MA, USA); Histone H3 (1:2500; 4499S;Cell Signaling Technology Inc, MA, USA); and SLC7A11 (1:500; 12691S, Cell Signaling Technology Inc, MA, USA). Immune complexes were detected with goat coupled anti-rabbit or horse coupled anti-mouse IgG antibodies (Cell Signaling Technology Inc).

### Clonogenic assay and proliferation assay

Cells were seeded in triplicate at a concentration of 500 cells per well into 6-well plates. After 24 hr, cells were continuously treated with indicated compounds for about 12 days. Colonies were fixed with fixation solution (5 parts methanol + 1 part acetic acid) at room temperature for 20 min and then stained with a solution of 0.4% (w/v) Coomassie Brilliant Blue G (B0770-25G) in H_2_O for 2 hours. For proliferation assay, cells were seeded into 96-well plates at a cell density of 2,000 cells per well, incubated overnight and treated with indicated compounds. Cell viability measurement was carried out after indicated treatment period using CellTiter-Glo kit (Promega, Cat. G7571) according to the manufacturer’s protocol.

### NADPH measurement

For NADPH measurement, cells were seeded into a T-75 flask at a cell density of 2 million cells per flask treated with indicated compounds. Cells were re-plated every 4 days and maintained in indicated compounds. 48 hr before NADPH measurement, cells were seeded at a 6-well plate at density of 0.3 million cells per well and maintained in indicated compounds. NADPH measurement was carried out after treatment using NADPH Assay Kit (Abcam,ab186031), and normalized to protein concentration measurement carried out using Pierce™ BCA Protein Assay Kit (Thermo Scientific, Cat. 23227).

### GSH/GSSG measurement

For GSH/GSSG measurement, cells were seeded into a T-75 flask at a cell density of 2 million cells per flask treated with indicated compounds. Cells were re-plated every 4 days and maintained in indicated compounds. 48 hr before GSH/GSSG measurement, cells were seeded into 96-well plates at a cell density of 30,000 cells per well and maintained in indicated compounds. GSH and GSSG measurement was carried out after treatment using GSH/GSSG-Glo kit (Promega, Cat. V6611) according to the manufacturer’s protocol.

### Model of experimental metastasis using MDA-MB-231-luc cells

#### Tumor implantation by tail-vein injection

All animals were treated in accordance with the recommendations of the National Institutes of Health and approved by the University of Texas at Austin Committee on Animal Care. All animal procedures were performed according to protocols approved by the University of Texas at Austin Institutional Animal Care and Use Committee (IACUC). Tail vein injection was performed as previously described ^41^. Six to eight-week-old female immunocompromised NOD SCID gamma mice (Stock #: 005557, Jackson Laboratory, Bar Harbor, ME, USA) were injected with MDA-MB-231-Luc parental and KEAP1 KO clone#1 cells via the lateral tail vein using 29 gauge needles, and followed up for metastases burden. In brief, 1×10^6^ cells suspended in 200 µl DMEM were injected into the tail vein of each mouse on day 0. After tumors became established in the lung on day 1, mice were randomized and treated with either AZD0156 or AZD1390. Mice (6 to 7 per treatment group) received the following agents by oral gavage as specified by the experimental protocols: vehicle (0.5% w/v HPMC, 0.1% w/v Tween 80) or ATM inhibitors (AZD0156 or AZD1390) at 20 mg/kg, QD for 5 days a week. After 6 or 7 weeks, animals were sacrificed and examined macroscopically and microscopically for the presence of metastases.

#### Drug treatment and imaging

Tumor growth was monitored weekly using bioluminescence imaging. Briefly, mice were intraperitoneally (I.P.) injected with 100mg/kg luciferin (in 200 μL) with a 25G needle and imaged using the IVIS Lumina instrument in the procedure room. Before the imaging, mice were anesthetized by using isoflurane for approximately less than 15 minutes at a time. All mice were sacrificed using a CO2 chamber and tissues were collected. Upon collection, tissues were fixed in 10% normal buffered formalin followed by sections and H&E immunostaining.

### Model of lung cancer xenograft using NCI-H1299-luc cells

All animals were treated in accordance with the recommendations of the National Institutes of Health and approved by the University of Texas at Austin Committee on Animal Care. All animal procedures were performed according to protocols approved by the University of Texas at Austin IACUC. On day 0, six to eight-week-old female and male mice were injected subcutaneously on the right flank with NCI-H1299 (2 × 10^6^ cells in 100 μL of RPMI 1640 with 50% Matrigel). After tumors became established on day7, mice (7 per treatment group) were randomized based on tumor volume and received the following agents as specified by the experimental protocols: vehicle (0.5% w/v HPMC, 0.1% w/v Tween 80) or AZD1390 (20 mg/kg) QD for 5 days by oral gavage. Animals were observed daily, and tumor volume and body weight were measured three times per week. Tumor volumes were calculated using the following formula: (length × width^2^)/2. In addition, tumor growth was monitored weekly using bioluminescence imaging, as previously described. During the procedures, any animal experiencing rapid weight loss (greater than 20%, measured weekly), debilitating diarrhea, rough coat, hunched posture, labored breathing, lethargy, persistent recumbence, jaundice, anemia, significantly abnormal neurological signs, bleeding from any orifice, self-induced trauma, impaired mobility, or has difficulty obtaining food or water was immediately euthanized. In addition, mice were euthanized when they became moribund. At week 4 after innoculation, all mice were sacrificed using a CO2 chamber and tissues were collected. Upon collection, tumor tissues were fixed in 10% normal buffered formalin and weighed followed by sections for H&E immunostaining. No human participant was involved in this study. All methods are reported following ARRIVE guidelines for the reporting of animal experiments.

### TCGA pan-cancer dataset analysis

Mutation status and expression level data (RSEM, batch normalized from illumine HiSeq_RNASeqV2) of ATM and KEAP1 in 32 TCGA PanCancer Atlas datasets were downloaded from c-Bioportal^42,43^. Correlations of ATM and KEAP1 were analyzed using the all samples (n = 10,071), breast invasive carcinoma samples (n = 994) and lung adenocarcinoma (n = 503).

### Statistical analysis

Statistical analyses were performed using GraphPad Prism 9 software (GraphPad Software). A value of p < 0.05 was considered statistically significant. Data are presented as mean ± SD.

*All methods were carried out in accordance with relevant guidelines and regulations*.

## Acknowledgements

We thank Dr. William M. Bonner of the National Cancer Institute (NCI), and Dr. Kyle M. Miller of the University of Texas at Austin, for their thoughtful critiques and their help with reviewing this manuscript. We are indebted to Dr. Can Cenik of the University of Texas at Austin for his critiques on this manuscript. This work is supported by the Cancer Prevention and Research Institute of Texas grant (CPRIT RR190101), the Center for Cancer Research of the National Cancer Institute, and the Alfred P. Sloan Research Foundation (FG-2021-16433).

## Author Contributions Statement

H.L. and U.W. designed the experiments. H.L. Y.L. and U.W. performed the analyses. H.L. Y.L., C.W., H.J.B, M.J., S.W., Y.X., and MB performed the experiments. M.B. provided animal models and helped with the experiments. H.L., H.J.B and M.B. edited the manuscript. Y.L. and C.C. helped with the bioinformatics analyses. U.W. wrote the manuscript with input from all authors.

