## Supplementary information for "CRISPR metabolic screen identifies ATM and KEAP1 as targetable genetic vulnerabilities in solid tumors"

1, Developmental Therapeutics Branch, National Cancer Institute, National Institutes of Health, 37 Convent Drive, Bethesda, MD 20892, USA. 2, Department of Molecular Biosciences, The University of Texas at Austin, Austin, Texas 78712, USA. 3, Institute for Cellular and Molecular Biology, The University of Texas at Austin, Austin, Texas 78712, USA. 4, Drug Dynamics Institute, College of Pharmacy, The University of Texas at Austin, 1400 Barbara Jordan Blvd., Austin, TX 78723, USA.

*** Corresponding author:**

Developmental Therapeutics Branch

National Cancer Institute

National Institutes of Health

37 Convent Drive,

Bethesda, MD 20892, USA.

Supplementary Fig. 1

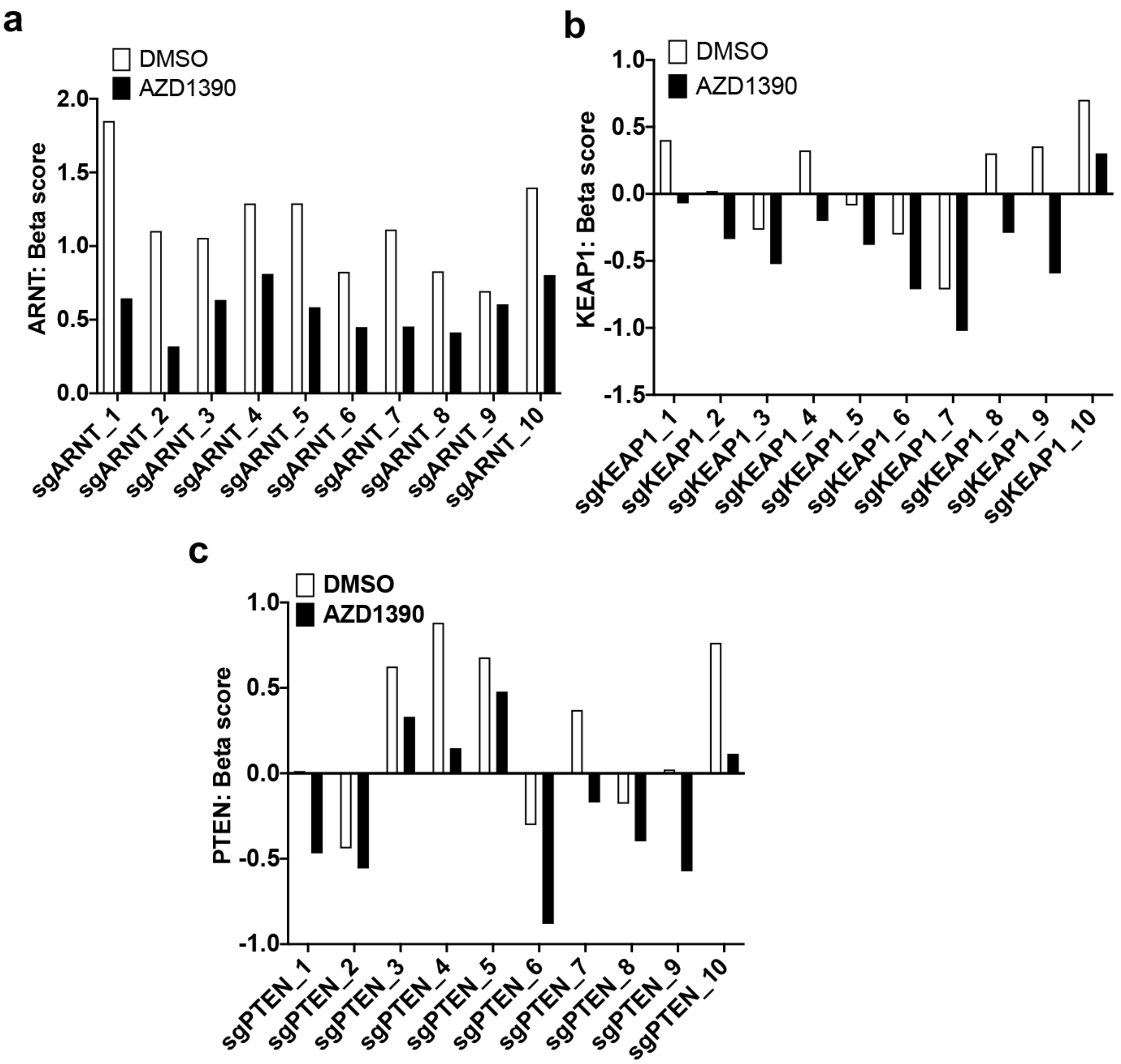


**Supplementary figure 1. Changes in abundance in the Metabolism-centered CRISPR/Cas9 knockout library screen of the individual sgRNAs of ARNT (a) KEAP1 (b) or PTEN (c) in presence or absence of ATM inhibitor (AZD1390).**

Supplementary Fig. 2

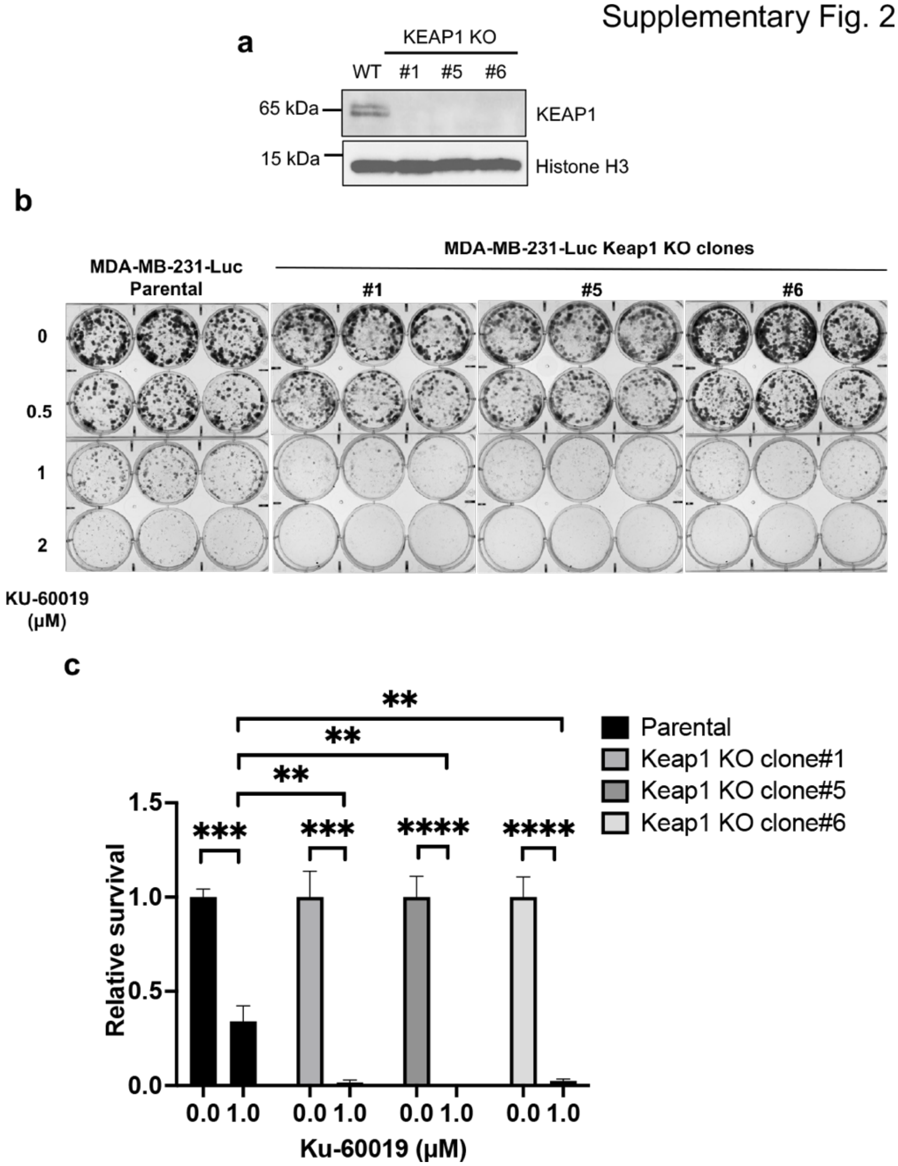


**Supplementary figure 2. Loss of KEAP1 sensitized MDA-MB-231 cells to ATM inhibition by KU-60019.** (a) Generation of *KEAP1* knockout MDA-MB-231-Luc cell lines by CRISPR/Cas9 approach. KEAP1 levels were probed by western blot. Histone H3 was used as a loading control. (b) Clonogenic assays of parental and *KEAP1* knockout cells (clones #1, #5, and #6) after treatment with ATM inhibitor (KU-60019) as indicated (0.5, 1, 2 μM). (c) Quantification of panel b. Statistical significance was determined by two-tailed unpaired student t test. Data are represented as mean ± SD; n = 3. **, $p < 0.01$; ***, $p < 0.001$.

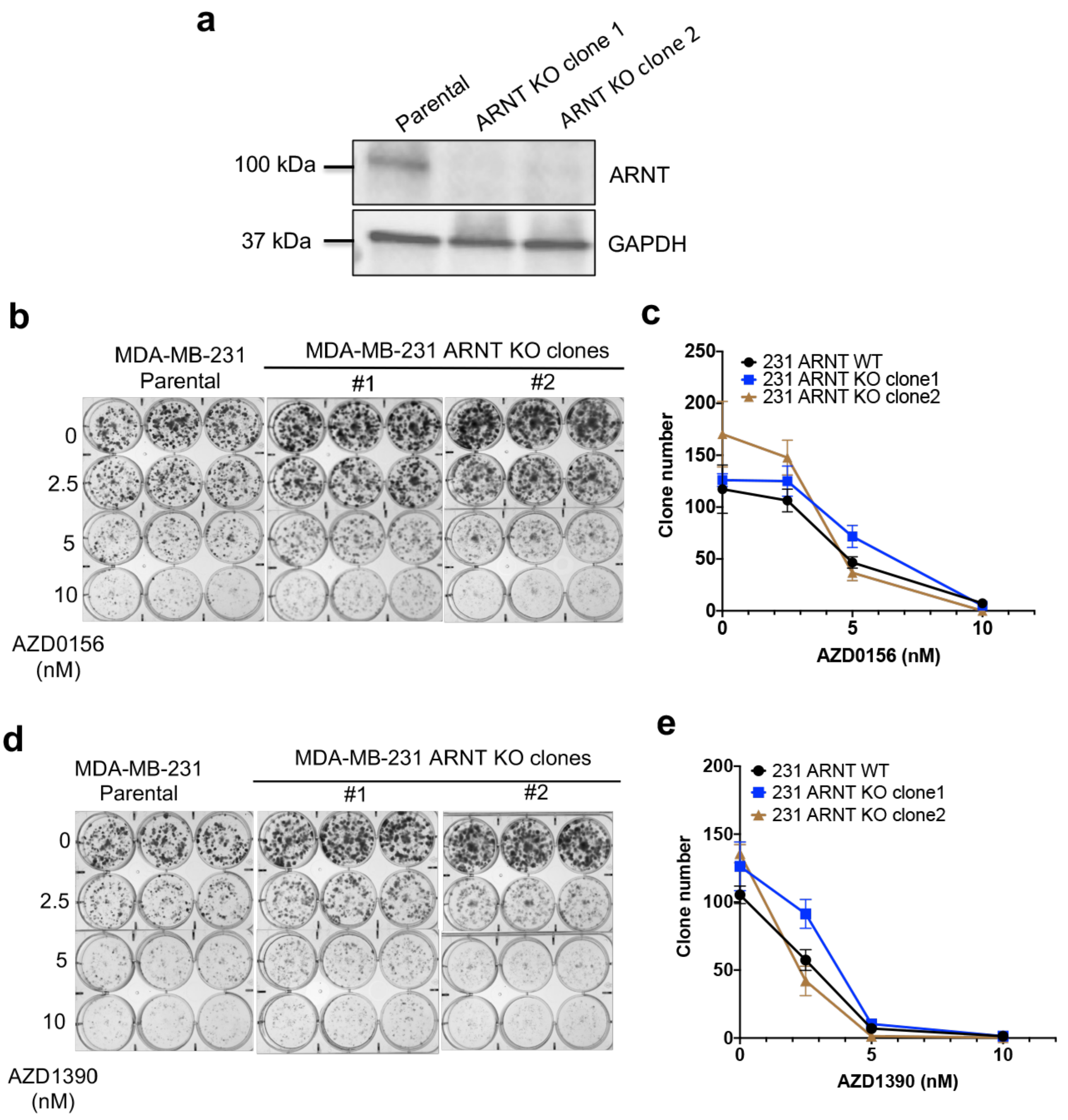


**Supplementary figure 3. ARNT loss did not sensitize MDA-MB-231 cells to ATM inhibition.** (a) Generation of ARNT knockout MDA-MB-231 cell lines by CRISPR/Cas9 approach. ARNT levels were probed by western blot. GAPDH was used as a loading control. (b) Clonogenic assays of parental and *ARNT*

knockout cells (clones #1 and#2) after treatment with ATM inhibitor (AZD0156) as indicated (2.5, 5, 10 nM). (c) Quantification of panel b. (d) Clonogenic assays of parental and *ARNT* knockout cells (clones #1, and #2) after treatment with ATM inhibitor (AZD1390) as indicated (2.5, 5, 10 nM). (e) Quantification of panel d.

Supplementary Fig. 4

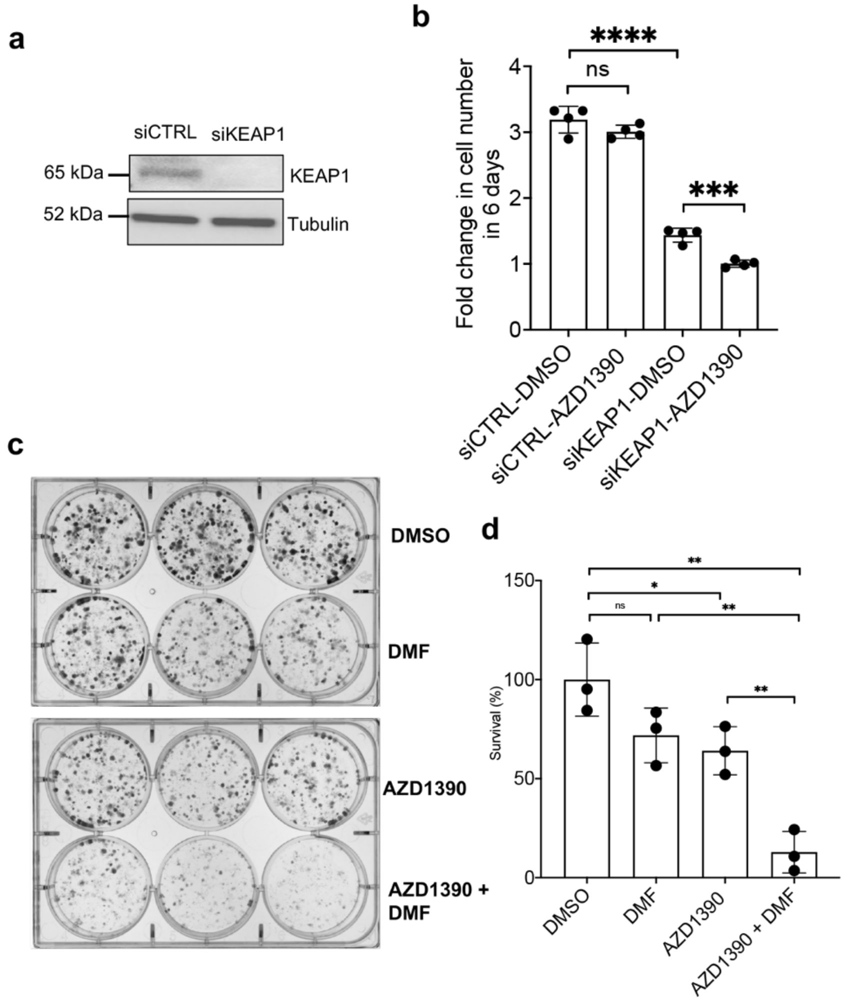


**Supplementary figure 4. KEAP1 deficiency or inhibition sensitized breast cancer cells to ATM inhibition.** (a) KEAP1 was knockdown in MCF7 cells using siRNA. KEAP1 levels were probed by western blot. Tubulin was used as a loading control. (b) Proliferation assays using MCF7 KEAP1 knockdown cells after treatment with ATM inhibitor (AZD1390) as indicated (50 nM). (c) Clonogenic assays using MDA-MB-231 cells after treatment with ATM inhibitor (AZD1390, 2.5 nM) and/or with KEAP1 inhibitor (Dimethyl fumarate, DMF) as indicated (10 μM). (d) Quantification of (c). Statistical significance was determined by two tail unpaired student t test. Data are represented as mean ± SD; n = 3. *, $p < 0.05$; **, $p < 0.01$; ***, $p < 0.001$; ****, $p < 0.0001$.

### Supplementary Fig. 5

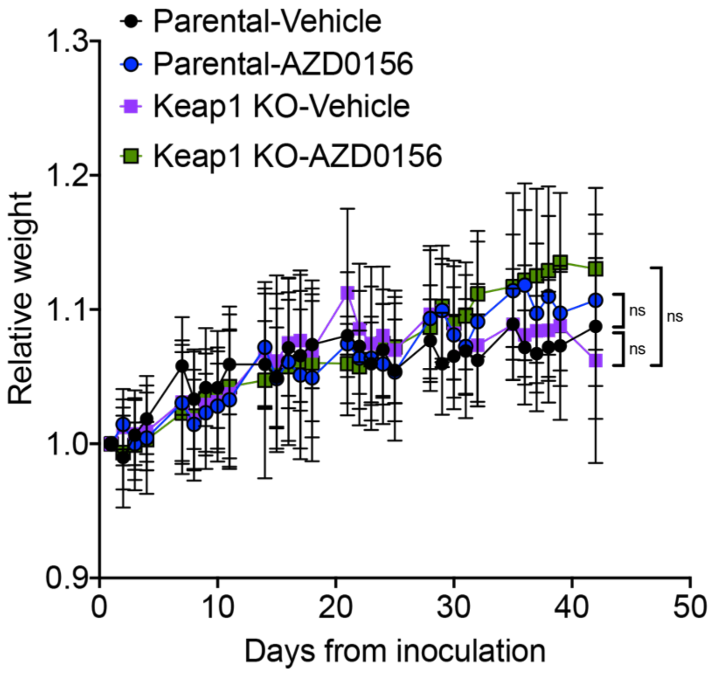


**Supplementary figure 5. Body weight of mice inoculated with MDA-MB-231-Luciferase parental or KEAP1 KO cells (clone 1) treated with either vehicle or AZD0156.** Statistical significance was determined by two-tail unpaired student t test. Data are represented as mean ± SD; n = 5 or 6. ns. not significant.
